# Inflammasome-mediated glucose limitation induces antibiotic tolerance in *Staphylococcus aureus*

**DOI:** 10.1101/2022.01.22.477360

**Authors:** Jenna E. Beam, Nikki J. Wagner, Kuan-Yi Lu, Sarah E. Rowe, Brian P. Conlon

## Abstract

*Staphylococcus aureus* is a leading human pathogen that frequently causes relapsing infections. Host-pathogen interactions have been shown to have substantial impacts on antibiotic susceptibility and the formation of antibiotic tolerant cells. In this study, we interrogate how a major *S. aureus* virulence factor, α-toxin, interacts with macrophages to alter the microenvironment of the pathogen, thereby influencing its susceptibility to antibiotics. We find α-toxin-mediated activation of the NLRP3 inflammasome induces antibiotic tolerance in the host cell cytoplasm. Induction of antibiotic tolerance is driven by increased glycolysis in the host cells, resulting in glucose limitation and ATP depletion in *S. aureus*. Additionally, inhibition of NLRP3 activation improves antibiotic efficacy in vitro and in vivo. Our findings identify interactions between *S. aureus* and the host that result in metabolic crosstalk that can determine the outcome of antimicrobial therapy.

## Introduction

Community-acquired methicillin-resistant *Staphylococcus aureus* (CA-MRSA) is the causative agent of multiple invasive infections, with high rates of morbidity and mortality (Cosgrove et al., 2003, Kourtis et al., 2019). In 2017, CA-MRSA sepsis contributed to over 20,000 patient deaths in the United States alone (Kourtis et al., 2019). Despite antibiotic therapy availability, treatment failure is common and often attributed to the formation of antibiotic tolerant cells (Kourtis et al., 2019, Labreche et al., 2013, Liu et al., 2020).

Antibiotic tolerant cells are a subpopulation of bacteria that enter a basal metabolic state, characterized by low levels of ATP (Rowe et al., 2020, Beam et al., 2021, Conlon et al., 2016, Huemer et al., 2021). In broth culture, glucose supplementation has been shown to resuscitate antibiotic tolerant cells by increasing their ATP levels (Conlon et al., 2016). Additionally, we have previously shown that reactive oxygen species (ROS) induce antibiotic tolerance via collapse of the tricarboxylic acid (TCA) cycle and ATP depletion (Rowe et al., 2020, Beam et al., 2021). The addition of exogenous glucose increased antibiotic susceptibility, even in the absence of a functional TCA cycle (Rowe et al., 2020). *S. aureus* virulence and proliferation in vivo is highly dependent on glucose, and its four glucose transporters, including 2 newly acquired and unique to *S. aureus*, demonstrate the importance of glucose acquisition to this pathogen (Vitko et al., 2015).

Due to the limitations of currently-approved antibiotics and a striking lack of new antibiotics in the pipeline, identifying and developing anti-virulence and/or host-directed therapeutics for the treatment of bacterial infections is becoming increasingly attractive (Fair and Tor, 2014, Beam et al., 2021, Cohen et al., 2018, Kane et al., 2018, Hua et al., 2015, Vu et al., 2020).

One of the major classes of virulence factors in MRSA are the pore-forming toxins, including leukocidins, phenol-soluble modulins, γ-hemolysin, and α-toxin. These toxins contribute to host cell death, initiate host cell signaling cascades, such as inflammasome activation, and mediate pathogen dissemination by facilitating escape from the host cell (Kebaier et al., 2012, Kitur et al., 2015, Craven et al., 2009, Cohen et al., 2018). Interestingly, antibody-mediated neutralization of α-toxin has been shown to improve infection outcome (Cohen et al., 2018, Vu et al., 2020, Hua et al., 2015, Ortines et al., 2018). However, how neutralization of α-toxin contributed to improved antibiotic efficacy was not determined.

α-toxin-mediated activation of the NOD-like receptor (NLR) pyrin domain-containing protein 3 (NLRP3) inflammasome contributes to *S. aureus* pathogenicity and immune evasion (Cohen et al., 2018, Craven et al., 2009, Liu et al., 2021b, Liu et al., 2021a). Once activated, the NLRP3 oligomerizes with itself and the apoptosis-associated speck-like protein containing a caspase recruitment domain (ASC) speck, forming the NLRP3 inflammasome. The NLRP3/ASC protein complex activates caspase-1, which cleaves pro-interleukin-1 beta (pro-IL-1β) and pro-IL-18 into mature IL-1β and IL-18, which are then secreted from the host cell. Secretion of IL-1β and IL-18 leads to increased inflammation and neutrophil recruitment to the site(s) of infection (Miller et al., 2007). The formation of gasdermin D pores downstream of NLRP3 activation can also result in inflammatory cell death, known as pyroptosis (Aachoui et al., 2013). Additionally, activation of NLRP3 has been shown to modulate host cell glycolysis (Finucane et al., 2019, Sanman et al., 2016, Shao et al., 2007). While the interaction between α-toxin and NLRP3 activation is well documented, the role of this interaction in antibiotic treatment outcome has not been determined.

In the current study, we aimed to determine if α-toxin-mediated activation of NLRP3 contributes to the formation of antibiotic tolerant *S. aureus* and if targeting activation of the NLRP3 signaling pathway is a potential host-directed therapeutic strategy that synergizes with antibiotic treatment.

## Results

### Loss of α-toxin increases antibiotic susceptibility

To determine the role of α-toxin in antibiotic tolerance, bone marrow-derived macrophages (BMDMs) and THP-1 human monocyte-derived macrophages (hMDMs) were infected with *Staphylococcus aureus* wildtype (WT) strain LAC or an α-toxin deletion mutant, Δ*hla*, followed by treatment with rifampicin (Fig 1AB, SFig1CD) or moxifloxacin (SFig 1A-E). Both rifampicin and moxifloxacin were chosen as these drugs are bactericidal and readily penetrate the macrophage by passive diffusion (Acocella et al., 1985, Barcia-Macay et al., 2006). At 24 hours post-infection (hpi), macrophages were lysed and CFU were enumerated. Compared to WT LAC, LAC Δ*hla* formed fewer antibiotic tolerant cells in the presence of both antibiotics (Fig 1, SFig 1).

**Figure 1.**
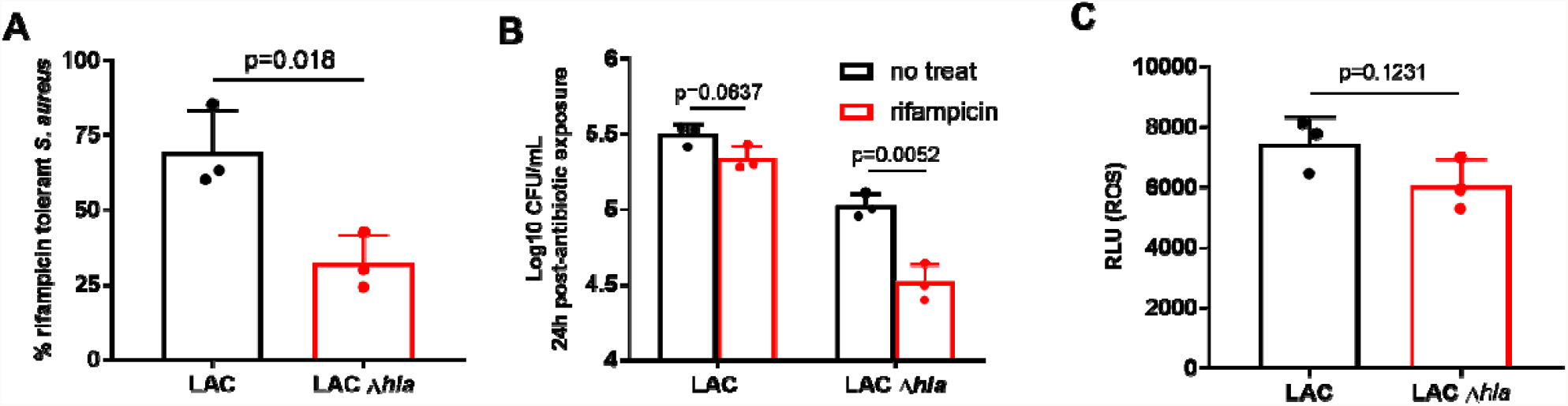
Loss of α-toxin increases antibiotic susceptibility in macrophages. (A,B) BMDMs were infected at MOI 10 for 45min, followed by treatment with 10μg/ml rifampicin for 24h. % survival (A) was extrapolated from CFU/ml (B). (C) ROS levels measured by L-012 luminescence in BMDMs infected for 45min at MOI 10 with LAC or LAC Δ*hla*. See also supplement figure 1. Statistical significance was determined by unpaired student’s t-test (A,C) or One-way ANOVA (B). Assays were performed in biological triplicate (n=3). Experiments are repeated a minimum of 3 times to ensure reproducibility. Bars represent mean + standard deviation.

We have previously shown that, in the phagolysosome, high levels of ROS, specifically peroxynitrite, induces an antibiotic tolerant state in *S. aureus* via collapse of central metabolism and reduced levels of ATP (Rowe et al., 2020, Beam et al., 2021). Given the high immunogenicity of α-toxin, we reasoned that perhaps when α-toxin is deleted, the macrophages would be less activated in the presence of the bacteria, leading to lower levels of ROS and thus decreased induction of antibiotic tolerant bacteria (Park et al., 1999). To measure ROS, BMDMs were infected with either WT LAC or LAC Δ*hla* for 1h, followed by addition of the ROS-sensitive luminescent probe L-012 or staining with fluorescein-boronate (FI-B; measures peroxynitrite) (Rios et al., 2016). Surprisingly, we observed no differences in ROS levels between WT or Δ*hla* infected macrophages (Fig 1C, SFig 1F).

### Inhibition of NLRP3 increases antibiotic susceptibility

Multiple studies have shown that α-toxin is a potent activator of the NLRP3 inflammasome (Craven et al., 2009, Cohen et al., 2018, Wang et al., 2020, Munoz-Planillo et al., 2013). Canonical NLRP3 activation is a two-signal process, where signal 1 is a priming step, typically toll-like receptor (TLR) signaling downstream of PAMP sensing. This leads to activation of NF-κb and upregulation of inactive NLRP3 monomers and pro-IL-1β and pro-IL-18. Upon receiving signal 2, NLRP3 becomes active and oligomerizes, which may lead to pyroptosis (Aachoui et al., 2013, Miller et al., 2007). Signal 2 can be a variety of stimuli, such as changes in calcium ion flux, mitochondrial damage, or, in the case of α-toxin, membrane pores that leads to potassium ion efflux (Craven et al., 2009, Cohen et al., 2018). To examine if NLRP3 activation contributes to the induction of antibiotic tolerance, we first measured caspase-1 activation and LDH secretion as proxies for NLRP3 signaling activation following infection with LAC or LAC Δ*hla*. BMDMs infected with WT LAC exhibited increased caspase-1 activation (Fig 2A) and LDH release (Fig 2B) compared to LAC Δ*hla* infected BMDMs. Next, we treated BMDMs with inhibitors of NLRP3 signaling, MCC950 or oridonin, prior to infection with LAC and treatment with rifampicin (Coll et al., 2015, Perera et al., 2018). Inhibition of NLRP3 increased rifampicin susceptibility in *S. aureus* (Fig 2C, SFig 2AB). Together, these data suggest that NLRP3 activation contributes to the induction of antibiotic tolerance and that inhibition of NLRP3 improves antibiotic efficacy in BMDMs.

**Figure 2.**
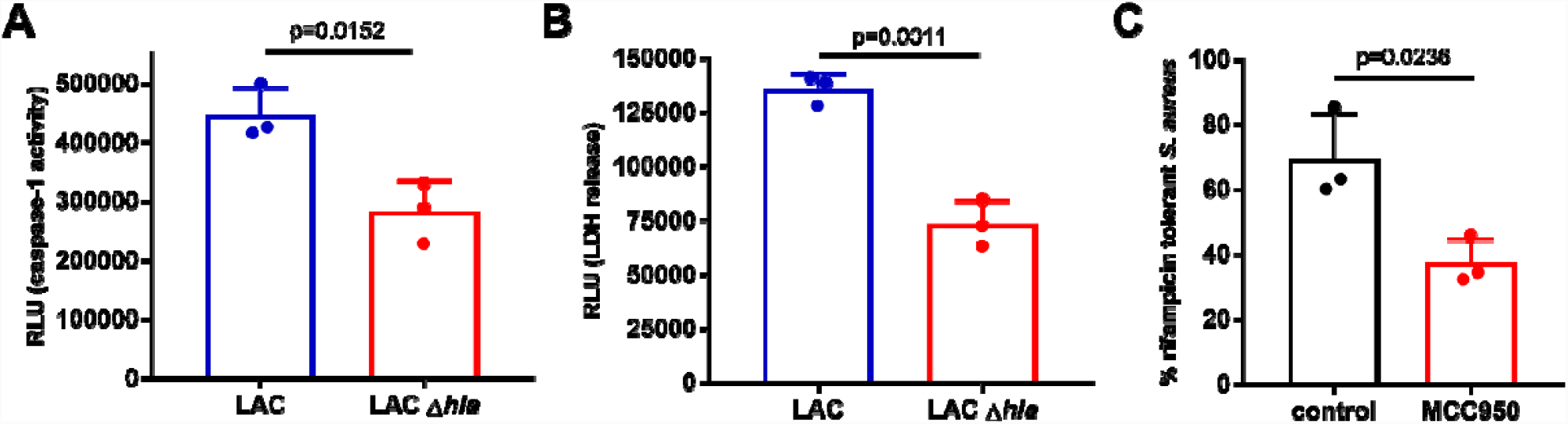
NLRP3 inhibition increases antibiotic susceptibility of *S. aureus*. (A) BMDMs were infected with LAC or LAC Δ*hla* for 24h followed by measurement of caspase-1 activity by luminescence. (B) BMDMs were left uninfected or infected with LAC or LAC Δ*hla* for 24h followed by quantification of LDH levels. (C) BMDMs were exposed to 100ng/ml lipopolysaccharide (LPS) for 2h, followed by replacement with serum-free media containing 10μM MCC950 for 30min prior to infection with WT LAC and treatment with 10μg/ml rifampicin for 24h. % survival of *S. aureus* recovered from BMDMs. % survival was extrapolated from CFU/ml at 24hpi (SFig 2A). Statistical significance was determined by unpaired student’s t-test. Infections were performed in biological triplicate (n=3). Experiments are repeated a minimum of 3 times to ensure reproducibility. Bars represent mean +standard deviation.

### α-toxin-mediated NLRP3 activation induces antibiotic tolerance in the host cytoplasm

Next, we aimed to determine how NLRP3 activation contributes to the induction of antibiotic tolerance. α-toxin has been shown to be important for phagosomal escape into the cytoplasm in non-professional phagocytes (Jarry et al., 2008). To determine if α-toxin is also important for phagosomal escape in macrophages, we performed confocal microscopy on J774A.1 macrophages infected with WT LAC or LAC Δ*hla* strains expressing GFP. By 24hpi, LAC Δ*hla* was still localized within the phagolysosome while the WT LAC was predominantly visible in the macrophage cytoplasm (Fig 3A,B). This data indicates that α-toxin is necessary for phagosomal escape in macrophages.

**Figure 3.**
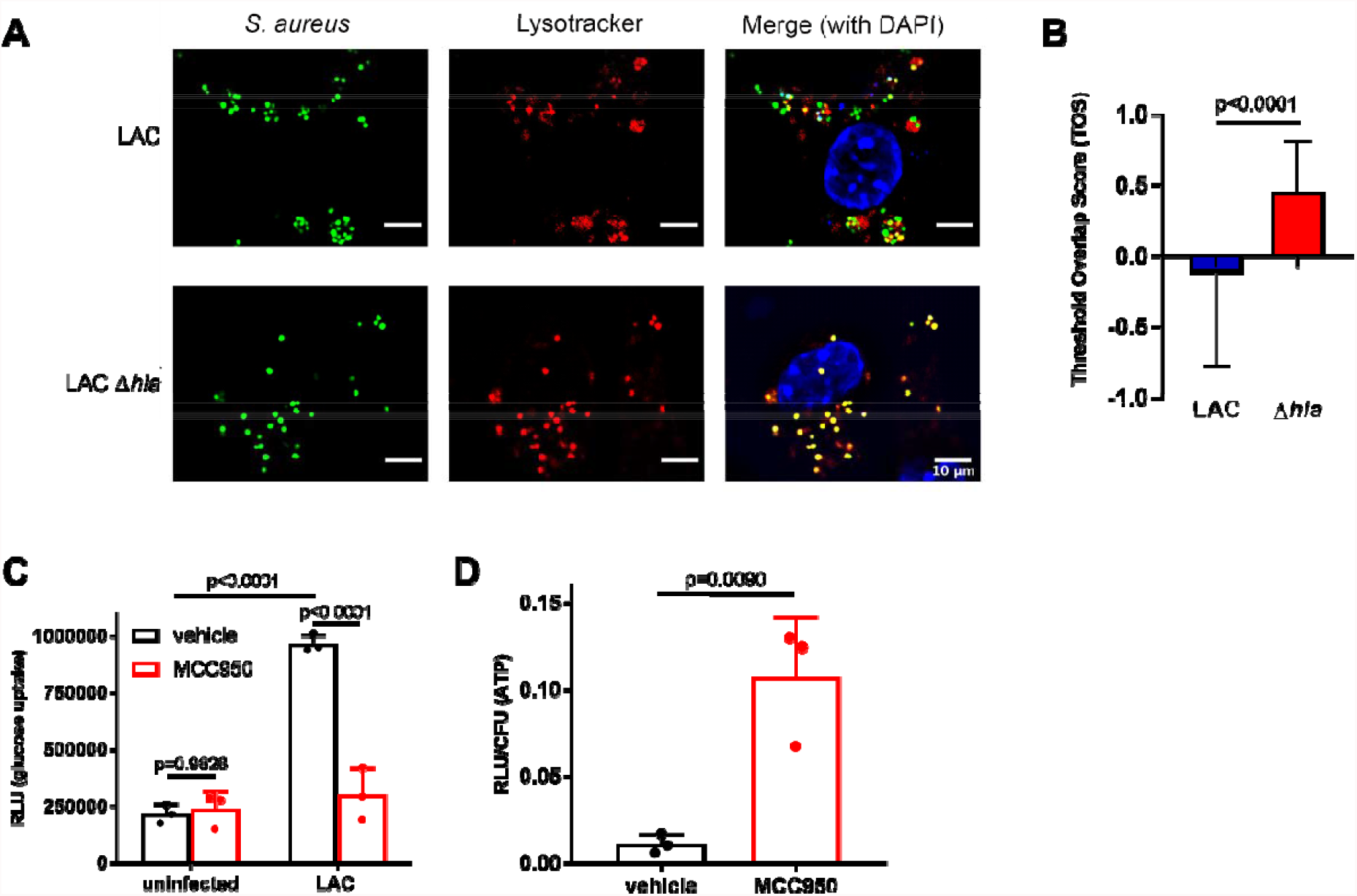
NLRP3 activation induces antibiotic tolerance in the macrophage cytoplasm. (A) Confocal microscopy of J774A.1 macrophages infected with GFP-expressing WT LAC or LAC Δ*hla* at 24h, followed by staining with LysoTracker (phagolysosome) and DAPI. (B) Total overlap score (TOS) indicates colocalization, where a TOS of 1 represents total colocalization and a TOS of −1 represents anti-colocalization. (C) Glucose uptake into BMDMs was measured at 24hpi using the Glucose Uptake-Glo assay kit (Promega). BMDMs were either untreated or treated for 2h with 100ng/ml LPS, followed by 10μM MCC950 and infection at MOI 10 with WT LAC. (D) ATP levels in *S. aureus* as measured by luminescence. BMDMs treated with and without MCC950 prior to infection at MOI 10 with LAC::*lux*. Luminescence was measured and normalized to CFU. Statistical significance was determined by Student’s t-test (B,D) or one-way ANOVA (C). All experiments were performed in biological triplicate (n=3) twice on two separate days (A-C) or three times on three separate days (D). Bars represent the mean + standard deviation.

TLR stimulation by bacterial PAMPS and NLRP3 activation leads to increased host cell glycolytic activity (Finucane et al., 2019, Sanman et al., 2016, Shao et al., 2007). Additionally, *S. aureus-*infected non-professional phagocytes have been shown to have decreased levels of intracellular glucose (Bravo-Santano et al., 2018). We reasoned that α-toxin-mediated NLRP3 activation leads to depletion of host cytoplasmic glucose, inducing antibiotic tolerance in *S. aureus* via nutrient deprivation. To test this, we measured glucose uptake into untreated or MCC950-treated BMDMs following 24h infection with WT LAC using the Glucose Uptake-Glo assay. After 24h, BMDMs were treated with 2-deoxyglucose (2DG), a glucose analog that is phosphorylated to 2-deoxyglucose-6-phosphate (2DG6P), but cannot be further metabolized by the host cell. Addition of glucose-6-phosphate dehydrogenase leads to reduction of NADP+ to NADPH, which converts proluciferin to luciferin. Relative light units (RLU) are therefore proportional to 2DG uptake into the host cells, which is indicative of host cell glycolytic activity. As shown in Figure 3C, BMDMs infected with *S. aureus* exhibit increased glycolytic activity, which is ameliorated by treatment with MCC950. These data indicate that inhibition of NLRP3 leads to decreased host cell glycolysis, which correlates with reduced antibiotic tolerant cells. Next, we wanted to measure ATP levels of LAC in untreated or MCC950-treated BMDMs. To measure ATP, LAC was transduced with a chromosomal *luxABDCE* cassette. The bioluminescent reaction is ATP-dependent and can thus be used as a proxy for bacterial ATP levels (Xu et al., 2014). BMDMs were infected with LAC::*lux* for 24h. BMDMs were then lysed and relative luminescence (RLU) was measured between the two strains. When NLRP3 was inhibited with MCC950, we observed increased ATP levels, which correlated with reduced to *S. aureus* antibiotic tolerant cells (Fig 3D).

To determine if the ability of *S. aureus* to run glycolysis correlates with changes in antibiotic tolerance, we infected untreated or MCC950-treated BMDMs with WT *S. aureus* strain JE2 or a glycolysis-deficient *pyk* transposon mutant. We hypothesized that in MCC950-treated BMDMs cytoplasmic glucose levels would be higher due to decreased host cell glycolysis (Fig 3C). If antibiotic tolerance is induced in the macrophage cytoplasm when *S. aureus* is starved of glucose, then a glycolysis-deficient mutant should still be tolerant to antibiotics regardless of cytoplasmic glucose availability (treatment with MCC950). Indeed, relative to the WT strain, the *pyk* mutant *S. aureus* remained tolerant, independent of cytoplasmic glucose availability, suggesting that the ability of *S. aureus* to catabolize glucose via glycolysis is directly proportional to the number of antibiotic tolerant cells (Fig 4A, SFig 3A).

**Figure 4.**
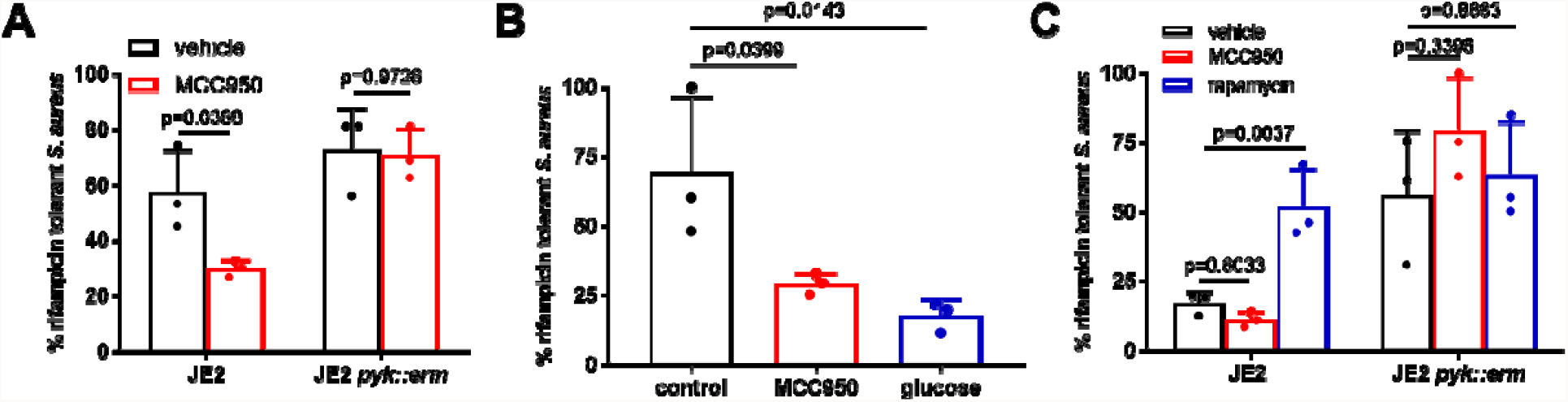
Glucose utilization is directly linked to antibiotic tolerance. (A) % rifampicin tolerance of *S. aureus* WT strain JE2 (black bars) or JE2 *pyk::erm* (blue bars) in BMDMs after 24h. BMDMs were infected at MOI 10 for 1h, followed by addition of 50μg/ml gentamicin +/-10μg/ml rifampicin. (B) % rifampicin tolerance of *S. aureus* WT LAC in untreated or MCC950-treated BMDMs. BMDMs were infected at MOI 10 for 1h, followed by addition of 50μg/ml gentamicin +/-10μg/ml rifampicin. At 20hpi, 0.2% glucose was added to the extracellular media, followed by CFU enumeration at 24h. (C) % rifampicin tolerance of *S. aureus* WT strain JE2 or JE2 *pyk::erm* in BMDMs after 24h. BMDMs were cultured in DMEM. Rapamycin-treated cells were incubated overnight in the presence of 100ng/ml rapamycin. BMDMs were infected at MOI 10 for 1h, followed by addition of 50μg/ml gentamicin +/-10μg/ml rifampicin. See also supplementary figure 3. Statistical significance was determined by One-Way ANOVA. All experiments were performed in biological triplicate (n=3) twice on two separate days. Bars represent mean + standard deviation.

Next, we wanted to determine if the addition of exogenous glucose could resuscitate and sensitize the cytoplasmic *S. aureus* antibiotic tolerant cells by stimulating *S. aureus* glycolysis. BMDMs were infected with WT *S. aureus* followed by treatment with or without rifampicin for 20h. At 20hpi, 0.2% glucose (∼0.01M) was added for 4h, at which point macrophages were lysed and CFU enumerated. Addition of glucose improved rifampicin susceptibility to a similar level observed with MCC950 treatment (Fig 4B and SFig 3B). This indicates that either blocking NLRP3-activation of host cell glycolysis or excess glucose is sufficient to sensitize antibiotic tolerant cells to rifampicin.

To further support the idea that glucose availability is a crucial determinant of antibiotic tolerance, we used rapamycin to repress glucose uptake by macrophages. Rapamycin selectively targets host cells but not *S. aureus*, thus allowing us to interrogate how the altered microenvironment affects the formation of antibiotic tolerant cells. To capture the effect of rapamycin on glucose limitation, infected BMDMs were cultured in a high-glucose medium (DMEM). In this scenario, we would expect fewer *S. aureus* antibiotic tolerant cells due to the excess amount of glucose (4.5g/L). Consistent with our hypothesis, there were increased *S. aureus* antibiotic tolerant cells in macrophages treated with rapamycin, highlighting the crucial role of glucose availability in antibiotic tolerance (Fig 4C, SFig 3CD). As expected, this effect cannot be readily detected in a low-glucose medium (MEM; SFig 3E). Altogether, these data suggest that α-toxin-mediated NLRP3 activation leads to increased host cell glycolysis, depleting cytosolic glucose levels, leading to reduced antibiotic tolerant cells as a result of nutrient deprivation following α-toxin-mediated phagosomal escape.

### NLRP3 inhibition improves antibiotic efficacy in murine bacteremia

To determine if NLRP3 inhibition improves antibiotic efficacy in vivo, we examined antibiotic treatment outcome in a systemic *S. aureus* infection on WT mice pre-treated with MCC950. Systemic infection was induced by tail vein intravenous (iv) injection, followed by treatment with rifampicin. Mice treated with MCC950 prior to infection and treated with rifampicin had statistically significantly lower bacterial burdens in their livers (Fig 5AB) and spleens (SFig 4) relative to vehicle control or rifampicin alone mice. These data suggest that NLRP3 inhibition improves antibiotic treatment efficacy against systemic *S. aureus* infection.

**Figure 5.**
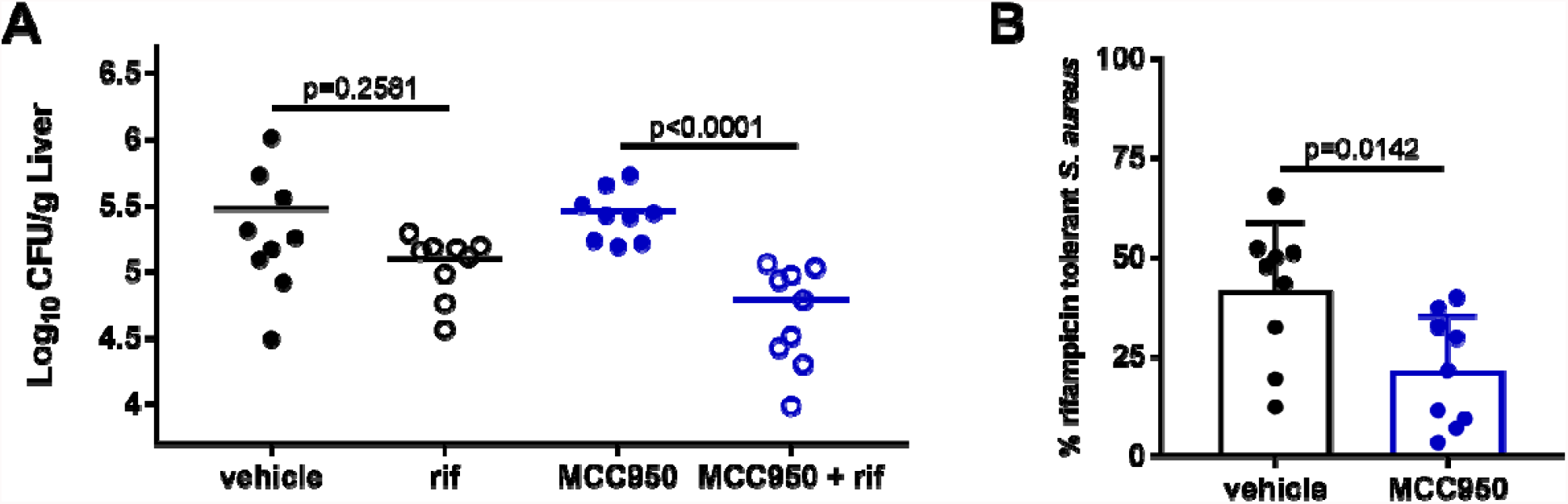
NLRP3 inhibition improves antibiotic efficacy against systemic *S. aureus* infection. WT C57B6/J mice were treated with 50mg/kg MCC950 by ip for 1 hour prior to infection, followed by tail vein iv infection with *S. aureus* strain HG003. At 24hpi, mice were administered 25mg/kg rifampicin (rif) or vehicle control by ip injection. (A) At 48hpi, *S. aureus* burden was enumerated in the liver. (B) % antibiotic tolerant *S. aureus* in vehicle versus MCC950-treated mice. See also supplemental figure 4. Each data represents one mouse from two experiments performed on two separate days (total n=9 per group). Statistical significance was determined by Mann-Whitney test comparing untreated to rifampicin treated.

## Discussion

*S. aureus* causes a variety of chronic and relapsing infections with high rates of antibiotic treatment failure, morbidity, and mortality. We have previously identified the intracellular niche as a potent driver of antibiotic tolerance in *S. aureus* (Rowe et al., 2020, Beam et al., 2021). Here, we find that inflammasome-mediated glucose limitation induces antibiotic tolerance in *S. aureus*.

NLRP3 activation is a two-signal process. Signal 1 is a priming step, typically TLR or other PRR recognition of PAMPs. Signal 2 can be a variety of different stimuli, including potassium ion efflux mediated by α-toxin, either directly or via packaging of *S. aureus* virulence factors in extracellular vesicles that are delivered to macrophages via endocytosis (Craven et al., 2009, Wang et al., 2020). TLR sensing of bacterial PAMPs, as well as NLRP3 activation, have been shown to shift macrophage to Warburg metabolism, characterized by increased glucose utilization and glycolytic flux (Shi et al., 2015, Finucane et al., 2019, Rother et al., 2019, Sanman et al., 2016, Shao et al., 2007). Additionally, α-toxin-mediated NLRP3 activation was recently shown to prevent immune clearance of *S. aureus* by recruiting mitochondria away from the phagolysosome, reducing mitochondrial ROS production and phagosomal acidification (Cohen et al., 2018). Other studies have shown that antibody neutralization of α-toxin during *S. aureus* pneumonia infection facilitates immune clearance and prolongs the antibiotic treatment window (Hua et al., 2015). However, how either NLRP3 activation, host cell metabolism, or neutralization of α-toxin impacts antibiotic efficacy has not been reported. Here, we show an intricate link between NLRP3 activation, host cell metabolism, and α-toxin wherein α-toxin activates NLRP3, increasing host cell glycolytic activity. Increased host cell glycolysis limits glucose availability for *S. aureus*, leading to cytoplasmic nutrient deprivation and subsequent tolerance following α-toxin-dependent phagosomal escape (Fig 6). By blocking NLRP3 activation, we are able to increase antibiotic susceptibility in *S. aureus* by stimulating *S. aureus* glycolysis.

**Figure 6.**
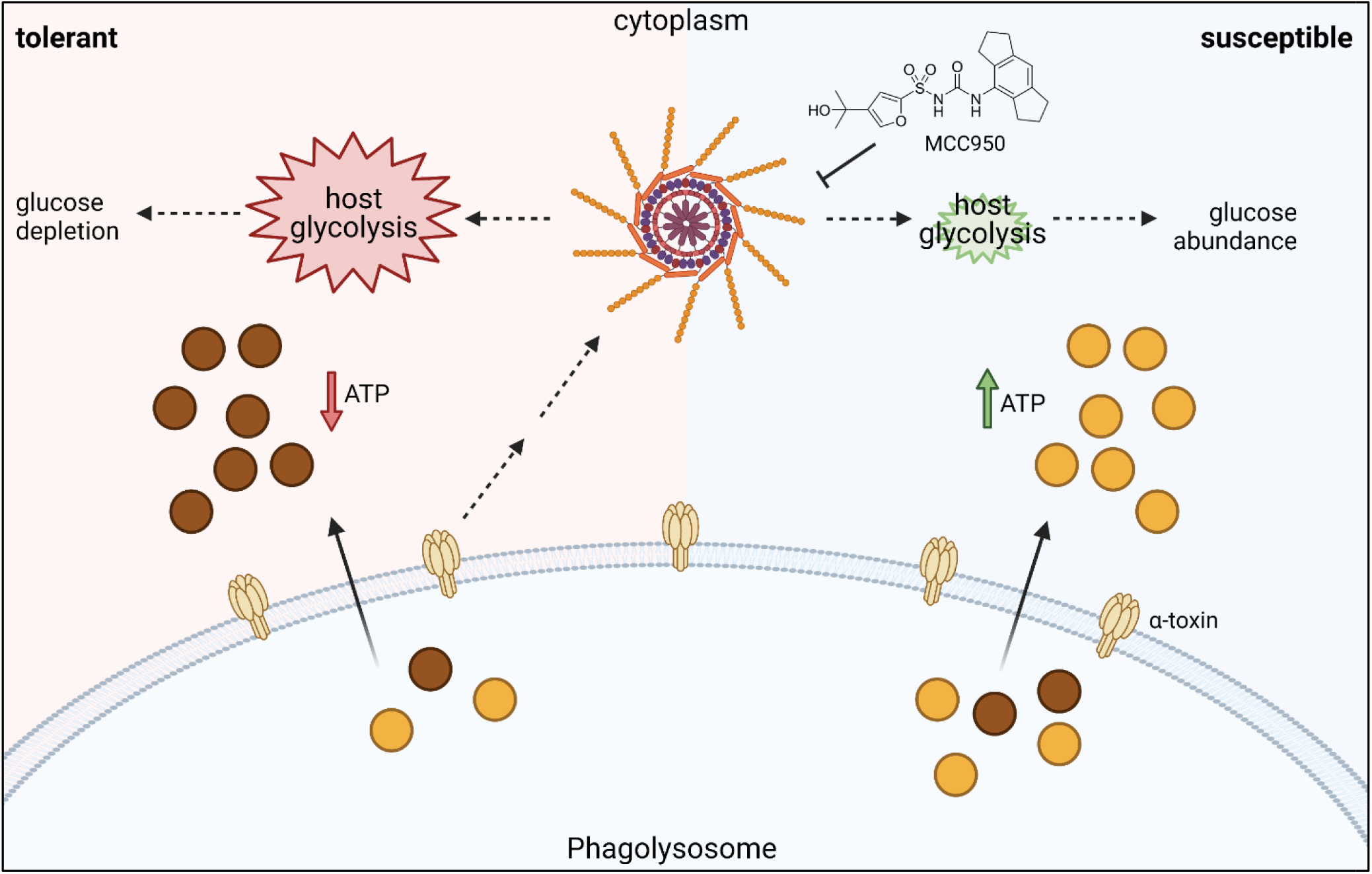
Proposed mechanism for NLRP3-mediated antibiotic tolerance in *S. aureus*. *Staphylococcus aureus* α-toxin activates NLRP3 and mediates escape from the phagolysosome. (left) Activation of NLRP3 increases host glycolytic activity, depleting the cytoplasm of glucose. *S. aureus* enters a low energy state (brown circles), hallmarked by low ATP, and is tolerant to antibiotics. (right) Pharmacologic inhibition of NLRP3 by MCC950 leads to decreased host glycolytic activity, leading to high levels of cytoplasmic glucose. *S. aureus* preferentially metabolizes glucose via glycolysis, leading to a high energy state (yellow circles), high ATP, and is thus sensitive to antibiotics.

The metabolic versatility of *S. aureus* greatly contributes to its success as a pathogen. As a facultative anaerobe, *S. aureus* is able to colonize and proliferate in a variety of host niches. As this and other studies demonstrate, the metabolic lifestyle of *S. aureus* in a given niche has significant impacts on antibiotic treatment efficacy, underpinning the importance of studying *S. aureus* antibiotic susceptibility in niche-specific contexts. The link between host cell metabolism and bacterial metabolism has previously been shown in other pathogens, including *Pseudomonas aeruginosa, Chlamydia trachomatis*, and *Mycobacterium tuberculosis (Mtb)*. A recent study showed that *P. aeruginosa* in the airway has adapted to utilize itaconate, a host-derived metabolite that accumulates during the proinflammatory response, as a nutrient source, leading to increased biofilm formation and chronicity of infection (Riquelme et al., 2020). *C. trachomatis* infection stimulates Warburg metabolism in infected cells, characterized by increased glycolysis and accumulation of nucleotides, facilitating *C. trachomatis* survival (Rother et al., 2019). Interestingly, Warburg metabolism was originally identified in tumor cells and is controlled by the tumor suppressor protein p53. Mutation of p53 in tumor cells leads to increased proliferation and inhibition of programmed cell death pathways (Vousden and Ryan, 2009). As activation of p53 inhibits host cell glycolysis, it reasons that acutely and reversibly targeting p53 during bacterial infection could improve antibiotic efficacy. In *Mtb* infected macrophages, interferon-γ-dependent hypoxia-inducible factor-1α (HIF-1α) causes a metabolic shift to aerobic glycolysis, which is essential for controlling *Mtb* infection (Braverman et al., 2016). HIF-1α is involved in a positive feedback loop that amplifies the proinflammatory immune response. Although a robust proinflammatory response was shown to be important for control of *Mtb* and *S. aureus* burden, it also leads to increased levels of reactive oxygen and nitrogen species (ROS/RNS). Work from our lab has shown that ROS potently induces antibiotic tolerance in *S. aureus* and nitric oxide has been shown to antagonize antibiotic killing of *Mtb - (Rowe et al., 2020, Beam et al., 2021, Liu et al., 2016)*, complicating the potential of targeting HIF-1α in the presence of antibiotics.

Overall, our results identify a complex signaling network whereby interactions between the *S. aureus* virulence factor α-toxin and the NLRP3 inflammasome result in metabolic crosstalk between host and pathogen that profoundly impacts antibiotic treatment efficacy.

## Materials and Methods

### Ethics Statement

All protocols used in this study were approved by the Institutional Animal Care and Use Committees at the University of North Carolina at Chapel Hill and met guidelines of the US National Institutes of Health for the humane care of animals.

### Bacterial Strains and Growth Conditions

*S. aureus* strains HG003, LAC (USA300), LAC::*luxABCDE*, LAC Δ*hla* (Nygaard et al., 2012), LAC Δ*hla* p*hla* (Nygaard et al., 2012) JE2, JE2 *pyk::erm* were routinely cultured in Mueller Hinton broth (MHB) at 37 °C and 225 r.p.m. Δ*hla* strains were grown in the presence of 250μg/ml spectinomycin and the complementation strain in 250μg/ml spectinomycin + 20μg/ml chloramphenicol. The transposon mutant JE2 *pyk::erm* was grown with 10μg/ml erythromycin, and LAC::*luxABCDE* in 10μg/ml chloramphenicol. LAC::*luxABCDE* was created via phage transduction of the *lux* cassette from JE2::*luxABCDE* (Liu et al., 2017).

### BMDM Isolation and Infection

Bone marrow from wildtype (WT) C57BL/6J mice (Jackson Labs) was isolated as described in (Amend et al., 2016). Bone marrow cells were differentiated for 7 days in Dulbecco’s Modified Eagle Medium (DMEM) + 10% FBS + L-glutamine + sodium pyruvate + sodium bicarbonate + 30% L929-conditioned media. After 7 days, cells were plated at 4×10^5^ cells/ml in minimum essential media (MEM) + 10% FBS + L-glutamine (complete MEM) or Dulbecco’s Modified Eagle Medium (DMEM) + 10% FBS + L-glutamine + non-essential amino acids + sodium pyruvate (complete DMEM) and allowed to adhere overnight at 37°C, 5% CO_2_. For assays with MCC950 and oridonin, BMDMs were primed for 2h with 100μg/ml lipopolysaccharide (LPS), followed by 30min treatment with 10μM MCC950 in serum-free media or 5 μM oridonin. Where indicated, BMDMs were treated with 100ng/ml rapamycin overnight. BMDMs were incubated with *S. aureus* LAC, LAC Δ*hla*, LAC Δ*hla* p*hla*, JE2, or JE2 *pyk::erm* at MOI 10 for 45min at 37°C, 5% CO_2_ to allow for internalization. Media was removed, cells were washed 1x with PBS, and media was replaced with complete MEM or DMEM as indicated + gentamicin 50μg/ml and/or rifampicin 10μg/ml and/or 50X MIC moxifloxacin as indicated (Peyrusson et al., 2020, Beam et al., 2021). For glucose sensitization experiments (Fig 4B), 0.2% (∼0.01M) glucose was added at 20hpi. At indicated timepoints, media was removed, cells were washed 3x with PBS and macrophages were lysed with 1% triton-x100. CFU were enumerated via dilution plating on tryptic soy agar (TSA) plates.

### THP-1 cell culture and infection

THP-1 monocyte-like cells were cultured in RPMI-1640 + 10% FBS + L-glutamine (complete RPMI). For differentiation into macrophages, THP-1 cells were seeded at 4×10^5^ cells/ml in complete RPMI + 20ng/ml phorbol 12-myristate 13-acetate (PMA) for 24h. After 24h, cells were weaned in complete MEM for 1h. Cells were infected as above, similarly to BMDM infection.

### ROS measurement

The luminescent probe L-012 (Wako Chemical Corporation) and fluorescein-boronate fluorescent (FI-B) probe were used to measure ROS. BMDMs were plated at 4×10^4^□cells per well in white tissue-culture-treated 96-well plates. For L-012, the cells were washed three times with PBS. L-012 was diluted to 150□µM in Hanks’ balanced salt solution (Gibco). Luminescence was read immediately using a Biotek Synergy H1 microplate reader. For FI-B, 25μM FI-B was added and fluorescence was read at 492nm/515nm (excitation/emission) using the plate reader as above. Data shown are representative of 3 independent assays of 3 biological replicates. Statistical significance was calculated using student’s unpaired t-test.

### Relative ATP measurement

*S. aureus* strain LAC::*luxABDCE* was used to infect BMDMs at MOI 10 as above. At indicated timepoints, BMDMs were washed and lysed as described above. Luminescence was read on Biotek Synergy H1 microplate reader. RLU were normalized to CFU.

### Glucose Uptake Assay

Untreated or MCC950-treated BMDMs were infected at MOI 10 for 1h with *S. aureus* LAC as above. After 1h, 50μg/ml gentamicin was added and cells were incubated for 24h. At 24h, glucose uptake was measured by Glucose Uptake-Glo Assay Kit (Promega) per manufacturer’s instructions.

### Caspase-1 Activity

Caspase-1 activity was measured in BMDMs infected with LAC or LAC Δ*hla* at MOI 10 as above. After 1h, 50μg/ml gentamicin was added and cells were incubated for 24h. At 24h, caspase-1 activity was measured using the Caspase-Glo 1 Inflammasome Assay kit (Promega) per manufacturer’s instructions.

### Microscopy Sample Preparation

J774A.1 cells were seeded at a density of 2×10^5^ per well on poly-L-lysine coated number 1.5 glass coverslips in 24 well plates. J774A.1 cells were propagated in complete DMEM and cultured for assays in complete MEM with 500ng/ml LPS. Cells were infected with either wild type LAC expressing GFP (LAC-GFP) (Kolaczkowska et al., 2015) or Δ*hla* LAC-GFP (this study) at an MOI of 10. Following infection, plates were spun at 1200xg for 2min. One hour post-infection (hpi), cells were washed 1x in PBS and media was replaced with MEM supplemented with lysostaphin 10µg/ml. One hour prior to harvest, lysotracker red (Invitrogen) was added to indicated samples at 100nM. At either 1 or 24 hpi times, cells were washed 3x with PBS and fixed with 4% paraformaldehyde at room temperature for 15 min. Fixed cells were washed 3x in PBS. DAPI was diluted to 2ug/ml in were incubated in PBS + 2% FBS. Samples were incubated with DAPI for 5 min. Coverslips were washed 3x in PBS and mounted on slides with ProLong Diamond (Life Technologies). Coverslips were sealed with nail polish before ProLong set to preserve the depth of the samples. Samples were imaged on a Zeiss LSM 700 Confocal Laser Scanning Microscope using a 63X/1.4 Plan Apo Oil objective lens and Zeiss ZEN 2011 software.

### Image Analysis

ImageJ and the plugin DeconvolutionLab2 (Sage, D. et al; Methods; 2017) were used to deconvolve the images. One slice in the middle of each Z stack was removed and analyzed for colocalization. EZcolocalization (Stauffer, W et al; Scientific Reports; 2018) was used to measures the overlap in signals above threshold intensity values, generating a Threshold Overlap Score (TOS) (Sheng, H. et al; Biol Open; 2016). The workflow for analysis was: Select one plane of the Z stack to analyze from each image. Open the red (Lysotracker) and green (S. *aureus*) channel images. Regions of interest (ROI) were selected based on the GFP signal of *S. aureus*. The EZcolocalization plugin was opened and set to analyze reporter 1 (red) and reporter 2 (green) with the selected ROI as the cell identification input. Thresholds were automatically determined using the “ Default” algorithm. TOS was calculated using the most intense 10% of pixels in each channel. For every analysis a metric matrix was generated to show the calculated values over multiple threshold combinations and visually check that the most appropriate threshold was used for analysis. TOS values were calculated for 295 wild type LAC-GFP and 424 Δ*hla* LAC-GFP. Statistical analysis was performed using an unpaired t-test using Graph Pad Prism 9 (version 9.3.1).

### Murine Bacteremia Model

WT C57BL/6J (Jackson #000664) mice were housed in a pathogen-specific free facility. For mouse infections, 8–10-week-old male and female mice were infected with ∼5×10^6^ CFU of *S. aureus* strain HG003 in 100μl PBS by intravenous (iv) injection. 1h prior to infection, mice were administered 50mg/kg MCC950 sodium in PBS (Selleck Chem #CP-456773) by intraperitoneal (ip) injection. Rifampicin was dissolved in vehicle (6.25% DMSO + 12.5% PEG300) at a final concentration of 6.25mg/ml. At 24hpi, mice were treated with 25mg/kg rifampicin or vehicle control by ip injection. At 48hpi, mice were euthanized via CO_2_ asphyxiation followed by cervical dislocation. Spleens and livers were harvested, homogenized, serially diluted, and plated on TSA plates for enumeration of bacterial CFU. Percent rifampicin tolerant cells was determined by comparing survivors after rifampicin treatment to survivors of the vehicle treated group. Wild type mice: vehicle n=9 and rifampicin n=9. The mean is indicated by a horizontal line. Statistical significance was calculated using the Kruskal Wallis One-Way ANOVA or the Mann-Whitney test as described in the figure legends. Blinding or randomization was not necessary as all outputs (CFU/g tissue) are objective.

## Author Contributions

B.P.C, and J.E.B. conceptualized the project; B.P.C, S.E.R., and J.E.B. wrote the manuscript; J.E.B, N.J.W., and K.L. performed the tissue-culture experiments; J.E.B. performed the animal experiments; J.E.B. and N.J.W produced figures; B.P.C. and S.E.R. provided funding for the project.

## Acknowledgements

This work was supported in part by NIH grants R01AI137273 to B.P.C and R03AI148822 to S.E.R. and a Burroughs Wellcome Fund investigator in the pathogenesis of infectious disease (PATH) award to B.P.C. The Microscopy Services Laboratory, Department of Pathology and Laboratory Medicine is supported in part by P30 CA016086 Cancer Center Core Support Grant to the UNC Lineberger Comprehensive Cancer Center. We thank Jovanka Voyitch for the α-toxin mutant strain, Janelle Arthur for equipment, and Mark Ross for assistance with animal infections. We thank Roger Plaut for sharing JE2-lux (SAP430). We thank Jenny Ting, Lance Thurlow, and Janelle Arthur for thoughtful discussions.

## Data availability

Additional data that support the findings of this study are available from the corresponding author, Brian P. Conlon, upon request.

