## Supplemental Figures for "Inflammasome-mediated glucose limitation induces antibiotic tolerance in *Staphylococcus aureus*"

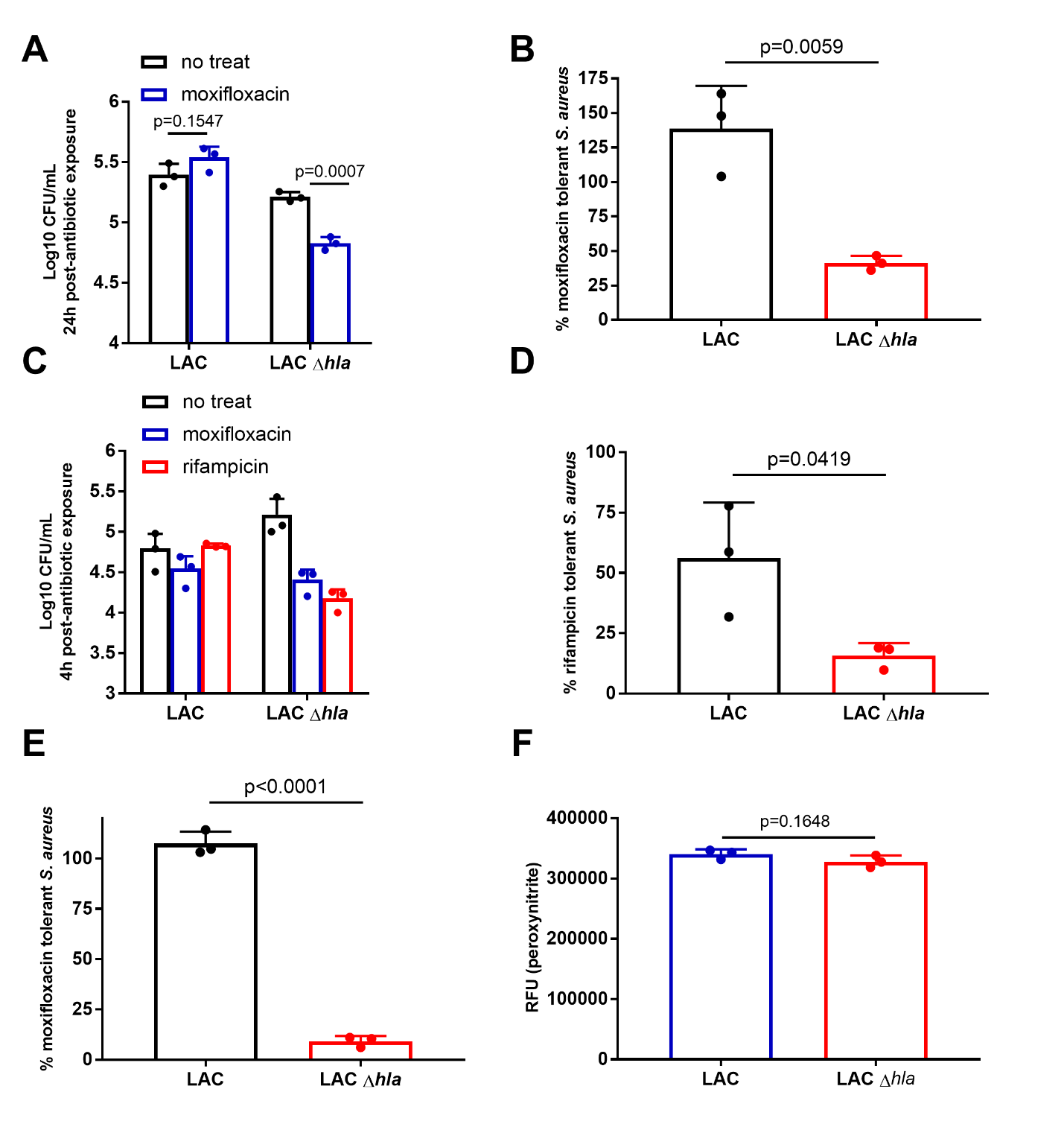

**SFig 1.** (A-B) Log CFU/ml (A) and % survival (B) of LAC and LAC *Δhla* infected BMDMs at 24hpi. BMDMs were infected at an MOI of 10, followed by treatment with 50X MIC (3μg/ml) moxifloxacin. (C) Log CFU/ml of LAC and LAC *Δhla* infected THP-1 monocyte-derived macrophages at 4hpi. THP-1 cells were infected at an MOI of 10, followed by treatment with 50X MIC (3μg/ml) moxifloxacin (blue) or 10μg/ml rifampicin (red). (D-E) % survival extrapolated from (C). (F) Peroxynitrite levels in BMDMs infected with LAC or LAC Δ*hla.* Peroxynitrite levels were measured by fluorescence. Bars represent mean + standard deviation.

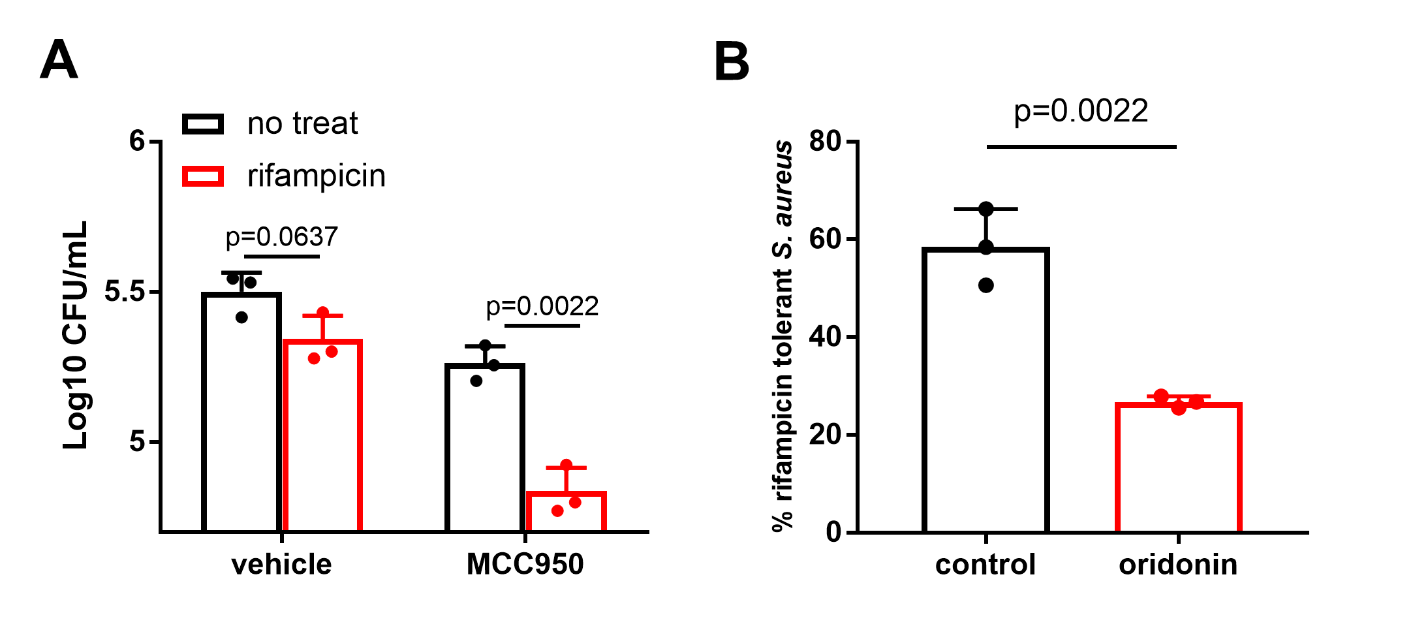

**SFig 2.** (A) Log CFU/ml of *S. aureus* strain LAC from untreated or MCC950-treated BMDMs +/- 10μg/ml rifampicin at 24h. (B) % survival of *S. aureus* strain LAC in BMDMs treated with 5uM oridonin. Symbols and bars represent mean +/- standard deviation.

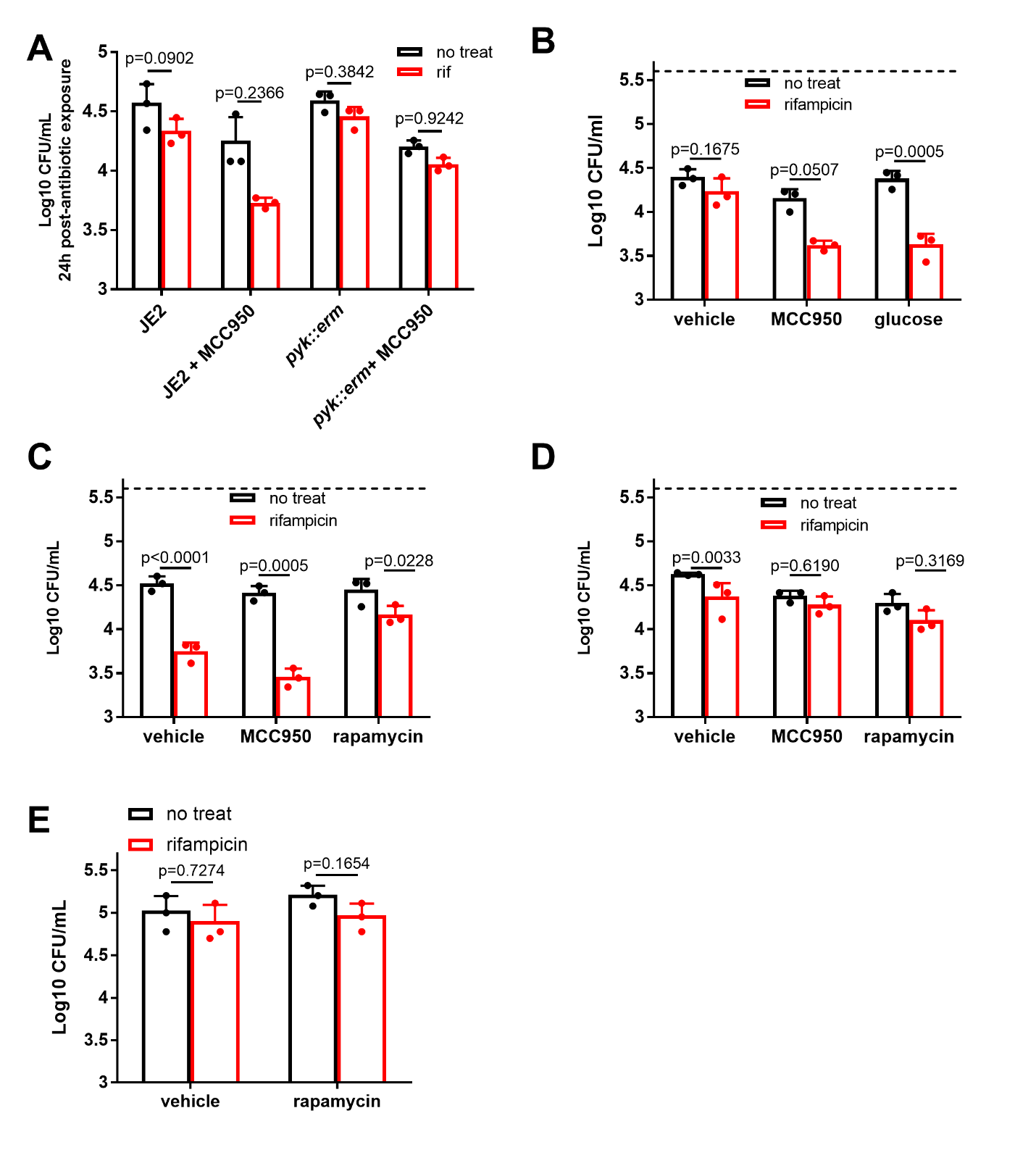

**SFig 3.** (A) CFU at 24hpi for JE2 and JE2 *pyk::erm* in BMDMs +/- MCC950. (B) CFU at 24hpi of LAC recovered from BMDMs treated with MCC950 or with the addition of exogenous glucose at 20hpi. (C-D) CFU for WT JE2 (C) and JE2 *pyk::erm* (D) at 24hpi recovered from BMDMs treated with MCC950 or rapamycin. Dashed black line represents average T0 CFU (B-D). (E) CFU at 24hpi for LAC recovered from BMDMs in MEM treated with 100ng/ml rapamycin, followed by treatment with 10μg/ml rifampicin. Bars represent mean +/- standard deviation.

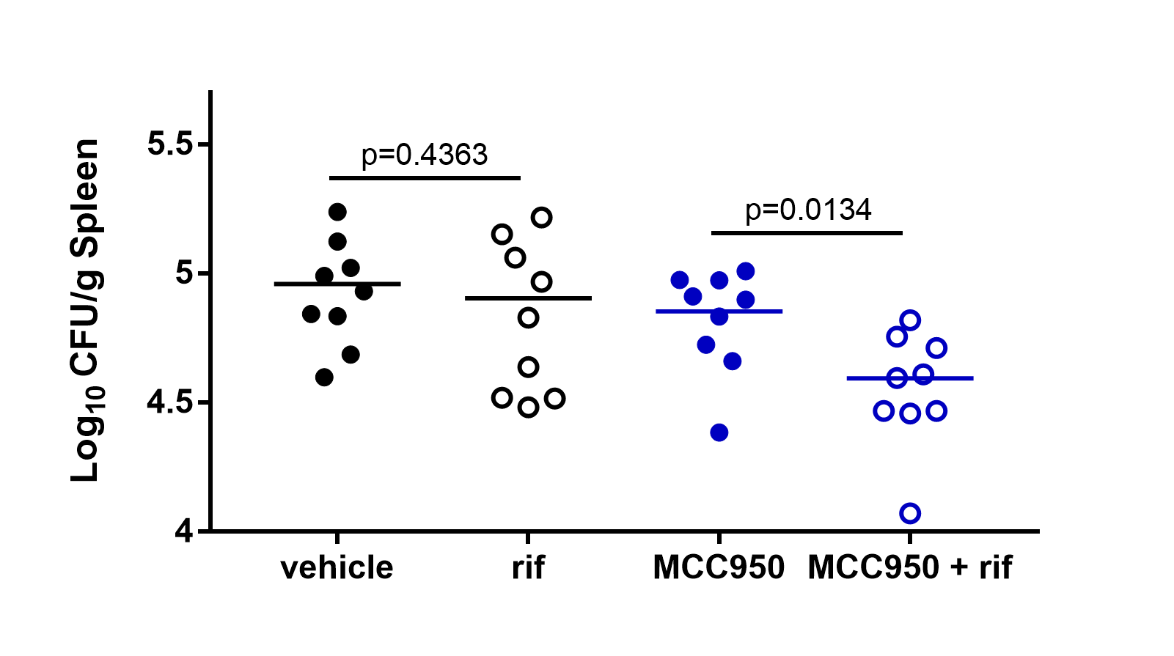

**SFig 4. NLRP3 inhibition improves antibiotic efficacy against systemic *S. aureus* infection.** WT C57B6/J mice were treated with 50mg/kg MCC950 by ip for 1 hour prior to infection, followed by tail vein iv infection with *S. aureus* strain HG003. At 24hpi, mice were administered 25mg/kg rifampicin (rif) or vehicle control by ip injection. (A) At 48hpi, *S. aureus* burden was enumerated in the spleen.
